# rTASSEL: an R interface to TASSEL for association mapping of complex traits

**DOI:** 10.1101/2020.07.21.209114

**Authors:** Brandon Monier, Terry M. Casstevens, Peter J. Bradbury, Edward S. Buckler

## Abstract

**Summary:** The need for efficient tools and applications for analyzing genomic diversity is essential for any genetics research program. One such tool, TASSEL (Trait Analysis by aSSociation, Evolution and Linkage), provides many core methods for genomic analyses. Despite its efficiency, TASSEL has limited means to use scripting languages for reproducible research and interacting with other analytical tools. Here we present an R package rTASSEL, a front-end to connect to a variety of highly used TASSEL methods and analytical tools. The goal of this package is to create a unified scripting workflow that exploits the analytical prowess of TASSEL in conjunction with R’s popular data handling and parsing capabilities without ever having the user to switch between these two environments. By implementing this workflow, we can achieve performances ranging from approximately 2 to 20 times faster than other widely used R packages for various functionalities.

**Availability and implementation:** rTASSEL is implemented in R using core TASSEL methods written in Java. The source code for rTASSEL can be found at https://bitbucket.org/bucklerlab/rtassel/src/master/. The source code for TASSEL can be found at https://bitbucket.org/tasseladmin/tassel-5-source/src/master/.

**Supplemental information:** The rTASSEL user manual and tutorials are freely available at https://maize-genetics.github.io/rTASSEL/. Supplemental material for this manuscript regarding performance metrics can be found in the attached file.

## Introduction

As breakthroughs in genotyping technologies allow for evermore available variant resources, methods and implementations to analyze complex traits are needed. One such resource is TASSEL (Trait Analysis by aSSociation, Evolution and Linkage). This software contains functionality for analyses in association studies, linkage disequilibrium (LD), kinship, dimensionality reduction, and genomic selection (Bradbury *et al*., 2007). While initially released in 2001, the fifth version, TASSEL 5, has been optimized for handling large data sets, and has added newer approaches to association analyses for many thousands of traits (Shabalin, 2012). Despite these improvements, interacting with TASSEL has been limited to either a graphical user interface with limited workflow reproducibility or a command line interface with a higher learning curve that can dissuade novice researchers (Zhang *et al*., 2009). To remediate this issue, we have created an R package, rTASSEL. This package interfaces the analytical power of TASSEL 5 with R’s data formats and intuitive function handling (R Core Team, 2020).

## Implementation

rTASSEL combines TASSEL’s abilities to store genotype data as half bytes, bitwise arithmetic for kinship analyses, genotype filtration, extensive forms of linear modeling, multithreading, and access to a range of native libraries while providing access to R’s scripting capabilities and commonly used Bioconductor classes (Figure S1) (Gentleman *et al*., 2004; Lawrence *et al*., 2013; Morgan *et al*., 2020). Since TASSEL is written in Java, a Java to R interface is implemented via the rJava package (Urbanek, 2019).

rTASSEL allows for the rapid import, analysis, visualization, and export of various genomic data structures (Figure S2). Diverse formats of genotypic information can be used as inputs for rTASSEL. These include variant call format (.vcf), HapMap (.hmp.txt), and Flapjack (.flpjk.*). Phenotype data can also be supplied as multiple formats including TASSEL formatted data sets or R data frame objects (Figure 1A).

**Figure 1.**
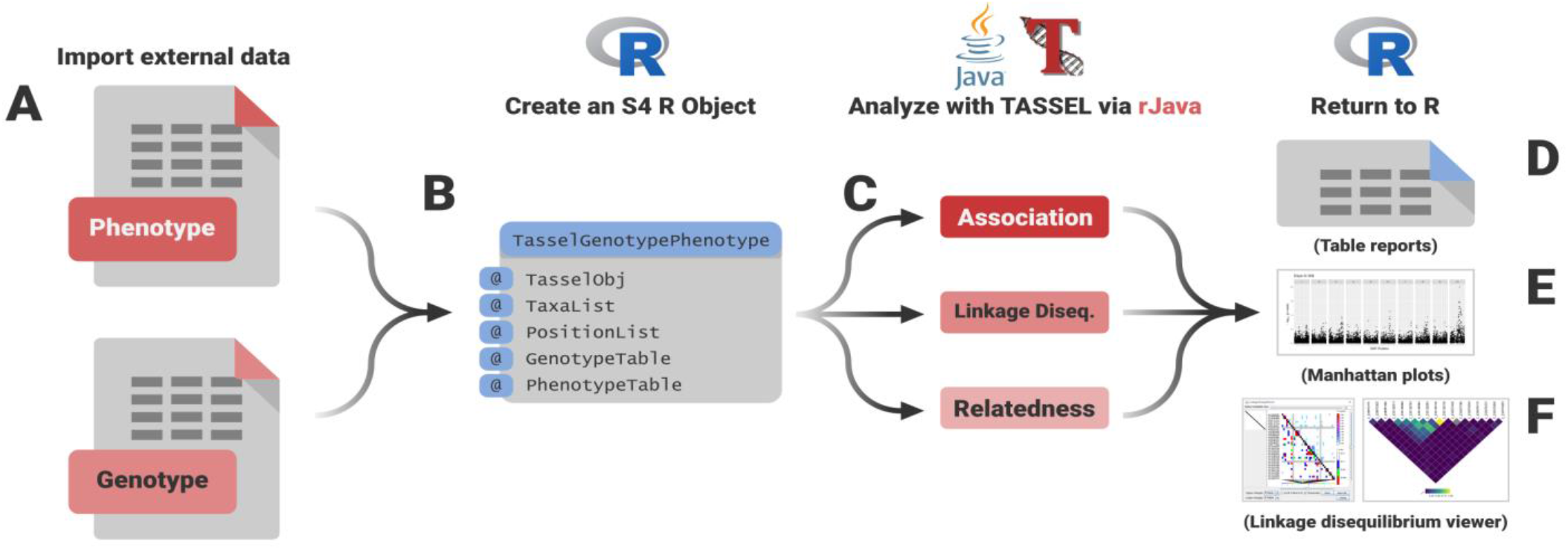
Genotypic and phenotypic data (A) are used to create an R S4 object (B). From this object, TASSEL functionalities can be called to run various association, linkage disequilibrium, and relatedness functions (C). Outputs from these TASSEL analyses are returned to the R environment as data frame objects (D), Manhattan plot visualizations (E) or interactive visualizations for linkage disequilibrium analysis (F).

Once data is imported, an S4 object is constructed that is used for all downstream analyses (Figure 1B, 1C). This S4 object contains slots that exclusively hold references to objects held in the Java virtual machine (JVM), which can be called with downstream analytical and filtration functions.

## Association methods

One of TASSEL’s most powerful functionalities is its capability of performing a variety of different association modeling techniques. rTASSEL allows for several types of association studies to be conducted by using one basic function with a variety of parameter inputs. This allows for implementing both least squares fixed effects general linear models (GLM) and mixed linear models (MLM) via the Q + K method (Yu *et al*., 2006) (Figures S3 and S4). If no genotypic data is provided to the GLM model, best linear unbiased estimates (BLUEs) can be calculated. Additionally, fast GLM approaches are implemented in rTASSEL which allow for the rapid analysis of many phenotypic traits and is approximately 4 times faster than the MatrixEQTL package (Figure S5) (Shabalin, 2012).

The data model for an analysis can be specified by a formula like R’s linear model function (R Core Team, 2020) which is shown as follows:

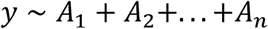

Where *y* is phenotype data and *A* is any TASSEL covariate or factor variables. This formula parameter along with several other parameters allow the user to run BLUE, GLM, or MLM modeling. Once association analysis is completed, TASSEL table reports of association statistics are generated as an R list which can then be exported as flat files or converted to data frames (Figure 1D). rTASSEL can also visualize association statistics which utilizes the graphical capabilities of the package, ggplot2 (Wickham, 2016) (Figure 1E).

## Other methods

rTASSEL can also be used for other analytical operations. For example, linkage disequilibrium (LD) can be estimated by the standardized disequilibrium coefficient, *D′*, as well as correlation between alleles at two loci (*r*^*2*^) and subsequent *P*-values via a two-sided Fisher’s Exact test. TASSEL Table reports for all pairwise comparisons and heatmap visualizations for each given metric can be generated via TASSEL’s legacy LD Java viewer or through R graphics (Figure 1F).

In order to perform MLM techniques, relatedness estimates (K) need to be calculated. TASSEL can efficiently compute this on large data sets by processing blocks of sites at a time using bitwise operations. This approach allows for approximately 2.5 times better performance compared to C methods in the statgenGWAS package (Fig S6) (Rossum and Kruijer, 2020). Several methods for calculating kinship in rTASSEL are implemented. By default, a “centered” identity by state (IBS) approach is used (Endelman and Jannink, 2012). Additionally, normalized IBS (Yang *et al*., 2011), dominance centered IBS (Muñoz *et al*., 2014), and dominance normalized IBS (Zhu *et al*., 2015) can be used.

rTASSEL can also be used for genomic selection by calculating genomic best linear unbiased predictors (gBLUPs). It proceeds by fitting a mixed model that uses kinship to capture covariance between taxa. The mixed model can calculate BLUPs for taxa that do not have phenotypes based on the phenotypes of lines with relationship information. When the analysis is run, the user is presented with the choice to run k-fold cross-validation. If cross-validation is selected, then the number of folds and the number of iterations can be entered. For each iteration and each fold within an iteration, the correlation between the observed and predicted values will be reported. If cross-validation is not selected, then the original observations, predicted values and prediction error variance (PEV) will be reported for all taxa in the dataset.

## Acknowledgements

This project is supported by the USDA-ARS, the Bill and Melinda Gates Foundation, and NSF IOS #1822330. We thank Sara J. Miller and Guillaume Ramstein for their insightful suggestions on this manuscript and pipeline testing.

