## Supplemental Methods and Figures for "rTASSEL: an R interface to TASSEL for association mapping of complex traits"

To achieve benchmarking results, several data sets were used. Genotypic and phenotypic maize data consisting of 279 samples, 3093 variant sites, and 1 measured trait were utilized for the analysis of variant call format (VCF) import, generalized linear model (GLM) association, mixed linear model (MLM) association, and kinship generation times (Flint-Garcia *et al.*, 2005). To illustrate the effectiveness of the fast association method, 100 simulated RNA expression traits for the prior genotype data was used. Trait data was generated using the `makeExampleDESeqDataSet()` function from the R package `DESeq2` (Love *et al.*, 2014). A larger genotypic data set consisting of 1,210 samples and 2,255,405 variant sites was also utilized for large VCF import and kinship generation times. All benchmarks were generated using the `microbenchmark()` function from the R package `microbenchmark` (Mersmann, 2019).

All benchmarks sans large VCF import and kinship generation times were evaluated 100 times and recorded on a workstation running 16 GB of RAM and 4 cores on an Intel® Core™ i5-6500 CPU with a clock speed of 3.20 GHz and. Large VCF import and kinship generation benchmarks were evaluated 10 times and recorded on a workstation running 256 GB of RAM and 12 cores on an Intel® Xeon® CPU E5-2643 v3 with a clock speed of 3.40GHz.

### Figures

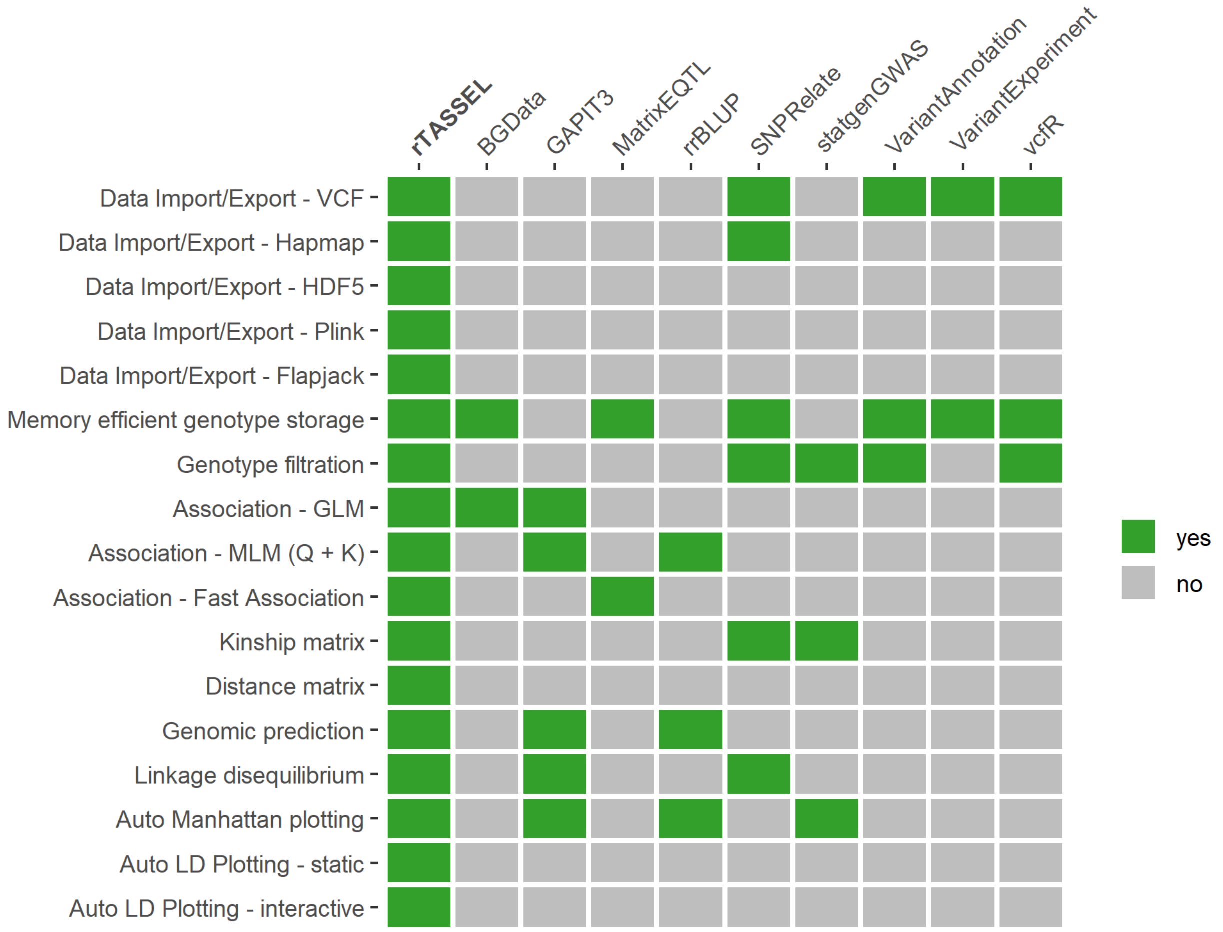


**Figure S1: Feature comparisons of rTASSEL with other R packages.** Features of rTASSEL (y-axis) are compared with other commonly-used R packages (x-axis). Packages that contain a specified feature are highlighted green (yes) and grey (no) if they do not contain a feature or are limited in scope. Association features for packages are based on if said package contains methods for generalized linear models, mixed linear models utilizing the "Q+K" method (Yu *et al.*, 2006), or multi trait fast association methods (Shabalin, 2012). Kinship and distance matrix features denote if a package can return an *n x n* matrix of values for further use. Packages that contain plotting features indicate if the package contains an automated plot feature instead of using base or grid-based R graphics (R Core Team, 2020) in conjunction with data output. The packages used for this comparison are BGData (Grueneberg and Campos, 2019), GAPIT3 (Wang and Zhang, 2020), MatrixEQTL (Shabalin, 2012), rrBLUP (Endelman, 2011), SNPRelate (Zheng *et al.*, 2012), statgenGWAS (Rossum and Kruijer, 2020), VariantAnnotation (Obenchain *et al.*, 2014), VariantExperiment (Liu *et al.*, 2020), and vcfR (Knaus and Grünwald, 2017).

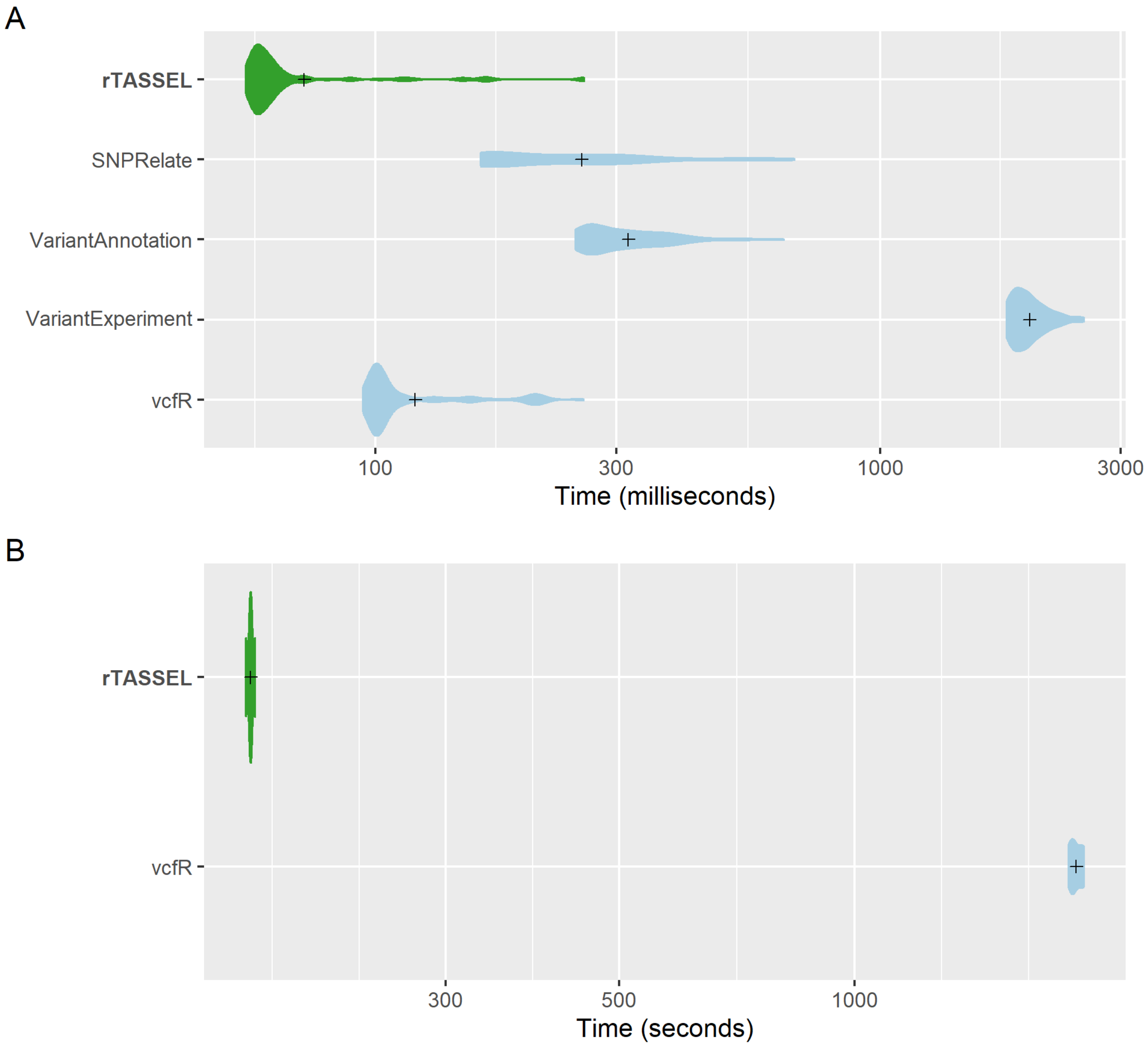


**Figure S2: VCF import time comparisons of genotypic data.** A distribution of replicated benchmark evaluations with recorded means (cross shapes) are plotted for rTASSEL and several R packages: SNPRelate (Zheng *et al.*, 2012), VariantAnnotation (Obenchain *et al.*, 2014), VariantExperiment (Liu *et al.*, 2020), and vcfR (Knaus and Grünwald, 2017). Import times are recorded for 279 samples x 3093 variant sites (A) and 1,210 samples x 2,255,405 variant sites (B).

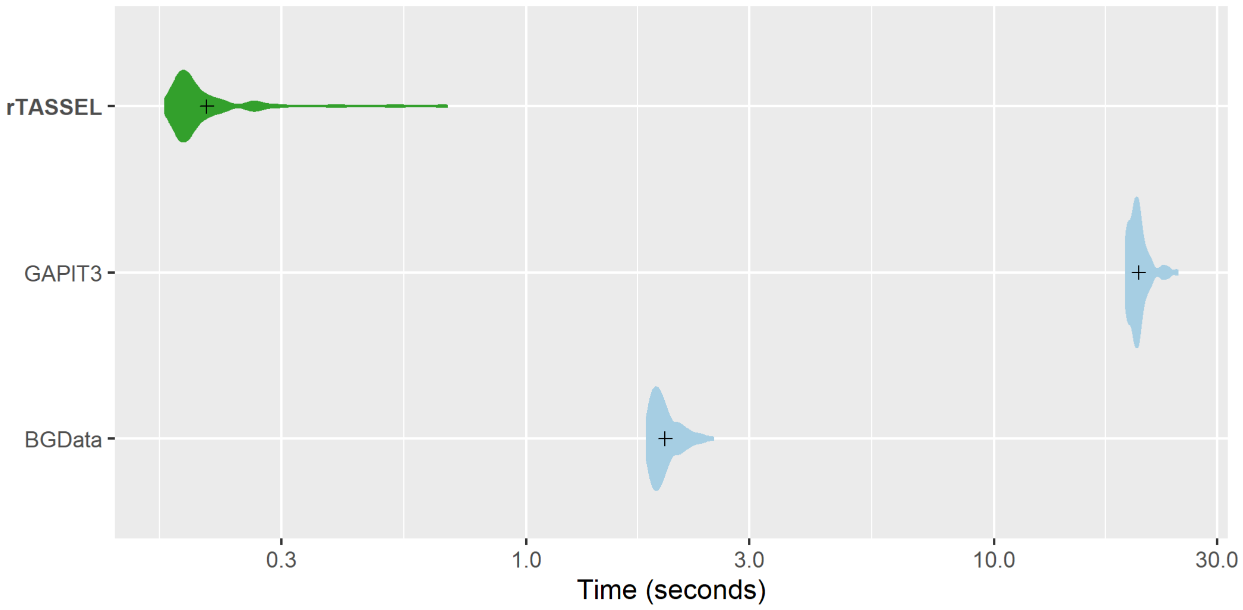


**Figure S3: GLM association time comparisons.** A distribution of replicated benchmark evaluations with recorded means (cross shapes) are plotted for rTASSEL and the R packages GAPIT3 (Wang and Zhang, 2020) and BGData (Grueneberg and Campos, 2019). Import times are recorded for 279 samples x 3093 variant sites and 1 measured trait.

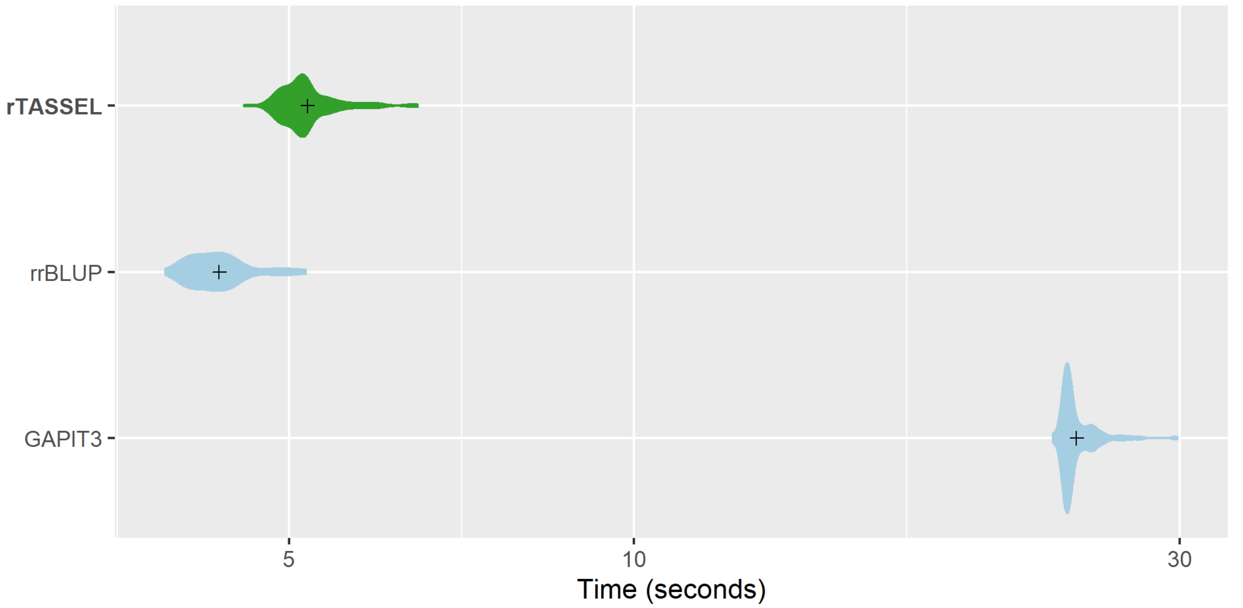


**Figure S4: MLM association time comparisons.** A distribution of replicated benchmark evaluations with recorded means (cross shapes) are plotted for rTASSEL and the R packages rrBLUP (Endelman, 2011) and GAPIT3 (Wang and Zhang, 2020). Import times are recorded for 279 samples x 3093 variant sites and 1 measured trait.

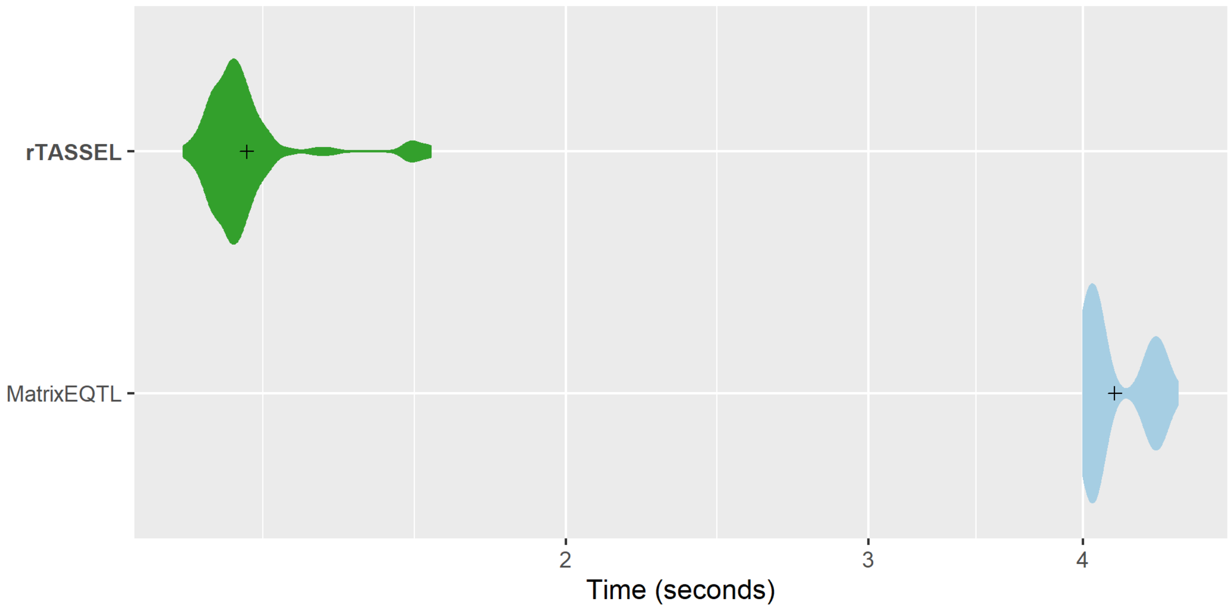


**Figure S5: Fast association time comparisons.** A distribution of replicated benchmark evaluations with recorded means (cross shapes) are plotted for rTASSEL and the R package MatrixEQTL (Shabalin, 2012). Import times are recorded for 279 samples x 3093 variant sites and 100 simulated RNA expression traits.

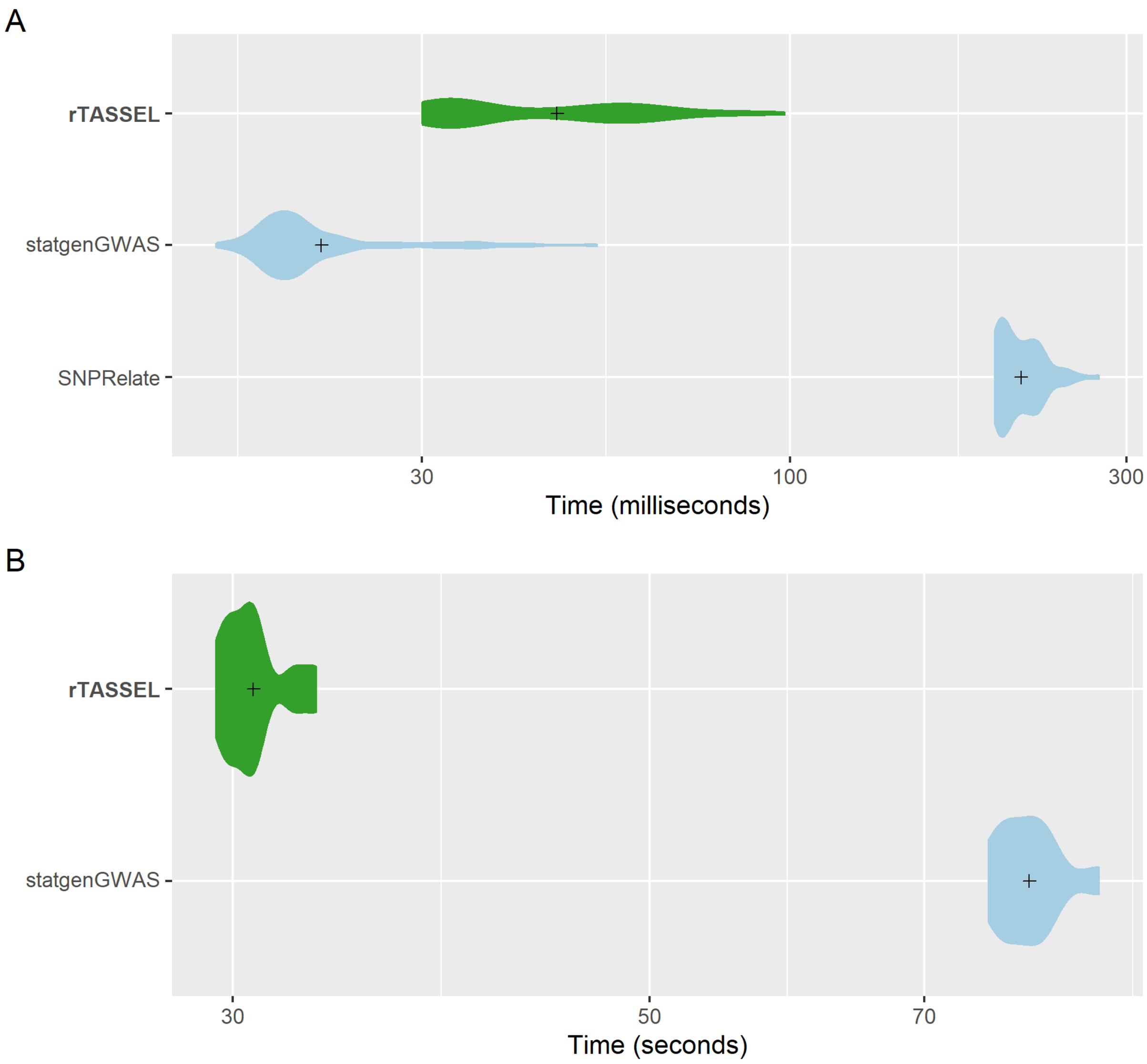


**Figure S6: Kinship matrix (IBS) generation time comparisons of genotypic data.** A distribution of replicated benchmark evaluations with recorded means (cross shapes) are plotted for rTASSEL and the R packages statgenGWAS (Rossum and Kruijer, 2020) and SNPRelate (Zheng *et al.*, 2012). Generation times are recorded for 279 samples x 3093 variant sites (A) and 1,210 samples x 2,255,405 variant sites (B).
